# Neural plasticity following surgical correction of strabismus in monkeys

**DOI:** 10.1101/319012

**Authors:** Mythri Pullela, Mehmet N. Ağaoğlu, Anand C. Joshi, Sevda Ağaoğlu, David K. Coats, Vallabh E. Das

**Affiliations:** College of Optometry, University of Houston, Houston, TX 77204; Department of Ophthalmology, Baylor College of Medicine, Houston, TX 77030

**Author notes:** **Corresponding author:** Vallabh E. Das, PhD, College of Optometry, University of Houston, 4901 Calhoun Rd, Houston, TX 77204. **Co-first authors:** Mythri Pullela and Mehmet N. Ağaoğlu are co-first authors.

**Keywords:** Strabismus, resection, recession, eye movements, non-human primate

## Abstract

The preferred treatment for correcting strabismus in humans involves the surgical manipulation of extraocular muscles (EOM). Although widely practiced, this treatment has varying levels of success and permanence, possibly due to adaptive responses within the brain or at the muscle. We investigated neural plasticity following strabismus surgery by recording responses from cells in the oculomotor and abducens nuclei before and after two monkeys with exotropia (divergent strabismus) underwent a strabismus correction surgery that involved weakening of the lateral rectus (LR) and strengthening of the medial rectus (MR) muscle of one eye. Eye movement and neuronal data were collected for a period of 6-10 months after surgery during a monocular viewing smooth-pursuit task. These data were fit with a first-order equation and resulting coefficients were used to estimate the population neuronal drive (ND) to each EOM of the viewing and deviated eyes. Surgery resulted in an ^~^70% reduction in strabismus angle in both animals that reverted towards pre-surgical misalignment by about 6 months after treatment. In the first month after surgery, the ND to the treated MR reduced in one animal and ND to the LR increased in the other animal, both indicating active neural plasticity that reduced the effectiveness of the treatment. Although these neuronal drive changes resolved by 6 months, we also found evidence for an inappropriate peripheral muscle adaptation that limited the effectiveness of surgery over the long term. Outcome of strabismus correction surgery could be improved by identifying ways to enhance ‘positive’ adaptation and limit ‘negative’ adaptation.

**Significance statement:** This is the first study of its kind to longitudinally follow behavioral and neural responses before and after a typical strabismus correction surgery in a monkey model for strabismus. We show the nature of muscle and neuronal plasticity that follows strabismus correction surgery.

## Introduction

Strabismus is a developmental disorder that affects ^~^4% of children worldwide (Von Noorden and Campos 2002, Govindan et al. 2005, Greenberg et al. 2007). In addition to ocular misalignment and sometimes visual acuity deficits (strabismic amblyopia), problems with binocular vision and oculomotor control such as deficits in processing disparity, poor stereo-acuity, disconjugate and cross-axis eye movements and nystagmus are also observed with strabismus and replicated in the monkey model (Lorenz 2002, Das 2008, Das 2009, Das 2016, Walton et al. 2017). The most common treatment strategy for strabismus is the surgical manipulation of specific extraocular muscles to re-align the eyes. Although popular and practiced widely, studies have shown that surgical approaches have varying levels of success and permanence, and often the patients tend to regain misalignment leading to multiple surgeries (Pineles et al. 2010, Yang et al. 2016). Ekdawi and colleagues suggested that the failure rate of surgical correction in children with intermittent exotropia rises to 84% by 15 years (Ekdawi et al. 2009). A more recent study by Chew et al. showed that post-operative success reduces from 75% at 1 week after treatment to 41% by the 5 year follow-up (Chew et al. 2016). Other forms of strabismus also have significant and variable failure rates (Louwagie et al. 2009, Habot-Wilner et al. 2012).

Fundamentally, the failure of surgery could be due to adaptive changes that are occurring at the periphery (muscle remodeling) or within central brain areas (central neural adaptation). For instance, when the globe was sutured to the orbit wall at an exotropic position in rhesus monkeys, sarcomere lengths were altered in the horizontal EOM, indicating that the muscle length has changed in response to the forced stretching of the sutured muscle (Scott 1994). Christiansen et al. studied the effects of resection surgery, a common strabismus correction technique to strengthen an apparently weakly acting muscle, performed on the lateral rectus in rats, and compensatory hypertrophy, i.e. increase in cell size, of the treated lateral rectus and the antagonist medial rectus was observed (Christiansen et al. 1988). Likewise, resection of rabbit EOM resulted in an increase in satellite cell activation in both the treated muscle and its antagonist muscle along with incorporation of new nuclei inside the myofibres (Christiansen and McLoon 2006), both thought to be indicators of active muscle remodeling.Antunes-Foschini and colleagues showed that the inferior oblique muscles obtained from strabismic patients contained more activated satellite cells than normal controls (Antunes-Foschini et al. 2006).

Previous investigations from our lab in monkey models for strabismus showed that innervation from the oculomotor nucleus to the extraocular muscles accounts for the state of strabismus and also cross-axis A/V patterns in animals who have not undergone any surgical manipulation (Joshi and Das 2011). Walton and colleagues studied the responses of neurons within the abducens nuclei in untreated strabismic monkeys and reported lower baseline firing rates than normal (Walton et al. 2014). Neurons within the supra-oculomotor area, which encodes vergence responses in normal animals, were found to encode horizontal misalignment in strabismic monkeys (Das 2012). More recent studies showed that electrical stimulation of the rostral superior colliculus causes a change in strabismus angle (Fleuriet et al. 2016) (Upadhyaya et al. 2017). Given the weight of evidence supporting a neural substrate for maintenance of strabismus, it is likely that neural plasticity following surgical manipulation of the EOM also contributes to the final strabismic state achieved after surgery. In this study, we investigated neuronal plasticity by comparing responses from motoneurons projecting to the horizontal recti before and after a typical resect-recess strabismus correction surgery. Our data shows evidence for significant neuronal plasticity that acts to reverse the intent of surgery, with larger changes observed to the resected MR muscle compared to the recessed LR muscle in one animal and the reverse in the other. Some of these data have been presented before in abstract form (Agaoglu et al. 2015, Pullela et al. 2015).

## Materials and Methods

### Subjects and rearing paradigms

All procedures in this study were performed according to National Institute of Health guidelines and the ARVO statement for the use of animals in ophthalmic and vision research and the protocols were reviewed and approved by the Institutional Animal Care and Use Committee (IACUC) at the University of Houston. The study consisted of two juvenile rhesus monkeys M1 and M2 (Macaca Mulatta) (^~^6 years of age, 9-10 kg). Strabismus was induced in infancy using an optical prism-rearing method. In this method, the infant monkeys wore a light-weight helmet fitted with a horizontally oriented Fresnel prism in front of one eye and a vertically oriented prism (20PD each) in front of the other eye. Prism-viewing starting from the first day of birth till they were ^~^4 months of age after which they were allowed unrestricted vision. Disruption of binocular vision (due to binocular decorrelation as a consequence of prism viewing) during the critical period is successful in inducing misalignment and other eye movement abnormalities associated with strabismus (Crawford et al. 1996, Crawford et al. 1996, Tusa et al. 2002, Das 2016, Walton et al. 2017).

### Surgical preparations and treatment of strabismus

When the monkeys were approximately 4-5 years of age, they underwent an aseptic surgical procedure while anesthetized with isoflurane (1.25-2.5%) to implant a titanium head stabilization post (Adams et al. 2007) to prevent head movements during experiments. In a second surgery, a scleral search coil was implanted in one eye using the technique of Judge and colleagues to measure eye movements (Judge et al. 1980). In this same surgery, we also implanted a 21mm diameter cylindrical titanium neural recording chamber centered at a stereotaxic location 1-mm anterior, 1-mm medial and 8-mm dorsal to stereotaxic zero for M1, and 3-mm anterior, 1-mm lateral, and 8mm dorsal to stereotaxic zero for M2. In both animals, the chambers were also tilted 20° dorsolateral to ventromedial in the coronal plane. This chamber location allowed access to both abducens and both oculomotor nuclei from within the same chamber. In a subsequent surgery, the fellow eye was also implanted with a search coil for binocular eye movement recording.

After initial behavioral training on standard oculomotor tasks and neurophysiological recording from each of the four motor nuclei to acquire pre-treatment data, the monkeys underwent a standard clinical resect-recess surgery (to correct the strabismus) that was performed by an expert strabismus surgeon and also one of the study authors (Figure 1). Muscle resection involves removing a section of muscle and therefore ‘strengthens’ the muscle due to its reduced length. Muscle recession involves repositioning the muscle insertion to a more posterior location and therefore ‘weakens’ the muscle because the muscle is less effective in transmitting torque to the globe. Since both monkeys were exotropic (exotropia or divergent strabismus is frequently attributed to weak medial rectus muscles and strong lateral rectus muscles), the medial rectus muscle of one eye was strengthened by resection, and the lateral rectus muscle of the same eye was weakened by recession. By design, surgical treatment was performed on only one eye (M1-left eye, M2-right eye) so that the fellow eye, including its EOM and corresponding motor nuclei, could serve as a control.

**FIGURE 1:**
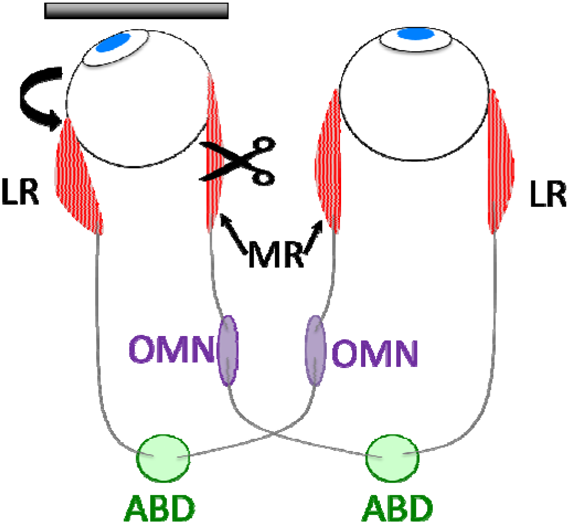
Schematic illustrating the surgical approach used to treat misalignment and the organization of motor nuclei projecting to each EOM. In strabismus surgery to correct exotropia, the ‘weak’ medial rectus (MR) is strengthened by shortening the muscle length via a resection procedure while the ‘strong’ lateral rectus (LR) is weakened by changing the point of insertion towards a more posterior location on the globe via a recession procedure. M1 underwent treatment on his left eye and M2 underwent surgery on his right eye. In this study, neuronal recordings were carried out within both oculomotor (OMN) and abducens (ABD) nuclei.

### Data Acquisition and Experimental Procedures

Binocular eye position was measured using the scleral search coil technique (Primelec Industries, Regensdorf, Switzerland). Calibration was performed as the monkey monocularly fixated within a ±2° window surrounding an optotype target that was back projected onto a tangent screen at a distance of 57cm. Visual targets were generated using a BITS# stimulus generation system (Cambridge Research Systems, UK) and presented using a DepthQ LCD projector (Lightspeed Design, Inc., Bellevue, WA, USA). Monocular viewing was enforced by occluding one of the eyes using liquid crystal shutter goggles (Citizen Fine Devices, Nagano, Japan) under computer control. Binocular eye position, target and neuronal data were collected as the monkeys performed a smooth pursuit task (0.3 Hz, ±15°) during monocular viewing with either the left or right eye. Eye and target position signals were passed through anti-aliasing filters at 400Hz before digitization at 2.79kHz with 12-bit precision (AlphaLab SNR system; Alpha-Omega Engineering, Nazareth, Israel). Raw spike data were collected at a sampling rate of 40kHz and sorted offline to generated time stamps of spiking activity (Spike 2 software, Cambridge Electronic Design). During further analysis using custom software developed in MATLAB (Mathworks, Natick, MA), spike time stamps were convolved with a 15ms standard deviation Gaussian to obtain a continuous spike density function of firing rate. Since the frequency of smooth-pursuit stimulation was low, target and eye movement data were further filtered using a Finite Impulse Response (FIR) low-pass filter with a cut-off of 20Hz or 50Hz.

### Data Analysis

A frequently used monocular first order equation (Eq 1) was used to fit the neuronal firing rates with smooth-pursuit eye movement data to estimate model coefficients, ‘K’, ‘R and ‘C’ (Sylvestre and Cullen 1999, Sylvestre and Cullen 2002, Joshi and Das 2011)):

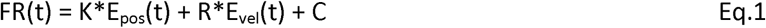

In this equation FR is firing rate of the neuron being recorded, E_pos_ and E_vel_ are the position and velocity of the eye that is controlled by the muscle to which the neuron projects; K is the position sensitivity of the neuron, R is the velocity sensitivity of the neuron and C is the baseline firing rate of the cell, i.e., firing rate when the eye that this neuron projects to is viewing a straight-ahead target. For example, the firing rate of a cell within the right OMN that shows increased burst-tonic activity for leftward movements (LTBT - projecting to the right eye medial rectus) would be modeled (Eq1) using the eye position and velocity of the right eye leading to estimates of ‘K’, ‘R’ and ‘C’ for that cell. Likewise, the activity of a LTBT cell within the left abducens nucleus (projecting to left eye lateral rectus) would be fit using the position and velocity information of the left eye. Some studies have suggested that a binocular model using position and velocity terms of both the ipsi- and contra-eye is a better representation of motoneuron responses (Zhou and King 1998, Sylvestre and Cullen 2002, Sylvestre et al. 2003). Since the main aim of the study was to investigate treatment effects, we decided that the monocular model to fit the neuronal firing and eye data would be the simplest and most interpretable approach.

Eye position signals were differentiated using a central difference algorithm, written in Matlab (Mathworks, Natick, MA) to obtain eye velocity. Previous studies have shown that position and velocity coefficients estimated during fast eye movements such as saccades are different from those estimated during slow eye movements such as smooth-pursuit or fixation (Sylvestre and Cullen 1999). Therefore, the smooth-pursuit data needed to be “de-saccaded” prior to fitting. Saccades were detected using a 40deg/sec velocity criterion, and eye and corresponding neuronal data during saccades were removed from the analysis. Model fitting was performed such that the eye position, velocity and neuronal data were resampled with replacement and fitted with the model in Eq. 1 and thereafter repeated 500 times. The model coefficients were deemed significant if the 95% confidence intervals for each coefficient did not overlap with zero. Eye movement and corresponding neuronal data from both right eye and left eye viewing conditions were concatenated during model fitting to develop the estimates of K, R and C.

After model coefficients were calculated, an estimate of the population neuronal drive (ND) to the lateral and medial rectus muscles of the deviated eye during monocular fixation with the fellow eye, was calculated using Eq. 2.

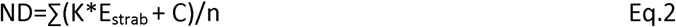

Where ND stands for neuronal drive, E_strab_ is the mean position of the non-viewing eye (position of deviated eye or strabismus angle) with the viewing eye looking straight ahead (i.e., at zero position), ‘K’ and ‘C’ are coefficients obtained from equation 1 for each of the cells projecting to the specific muscle in question, and ‘n’ is the number of cells recorded from the corresponding nucleus. Note that the coefficient R is not considered in Eq. 2 since ND is computed for a fixation condition (i.e., eye velocity is zero). In this framework, ‘ND’ is equivalent to the average population neuronal activity innervating the extraocular muscles (either LR or MR depending on the nucleus) of the eye under the cover during fixation. In a neural sense, these ‘ND’ are the reason that the eye under cover is deviated.

Neural data were collected longitudinally before and after treatment. For statistical analysis, the estimated measures were lumped and compared at three time intervals: ‘Pre’ - data recorded before the surgery, P1 - data recorded from day-1 to ^~^one month after treatment, P6 - data recorded from 6-10 months after treatment. Statistical testing was carried out using a one-way ANOVA at significance level of 0.05 following by Holm-Sidak method for post-hoc testing unless otherwise specified.

## Results

Prior to treatment, Monkey M1 had a mean exotropia of ^~^15° with the left eye viewing (LEV) and ^~^30° with the right eye viewing (REV) while M2 showed an exotropia of ^~^20° during REV and ^~^35° during LEV. On the first day following a resect-recess surgery on one eye, eye misalignment reduced by ^~^70% of pre-surgical values in both M1 and M2 when viewing with the treated eye. Figure 2 shows the longitudinal progression of strabismus angle in M1 and M2 before and after surgical correction. The data show that by the end of the P6 recording period, both monkeys showed large angle exotropia once again. Additional details of the longitudinal change in alignment and dynamics of eye movements can be found in our previous publication (Pullela et al. 2016). Note that the data points on the plot are chronologically arranged based on days when neural recording yielded cells. Frequently, more than one cell was recorded on a specific day. Alignment data were also recorded on days that did not yield successful neuronal recording but these are not shown in the plot.

**FIGURE 2:**
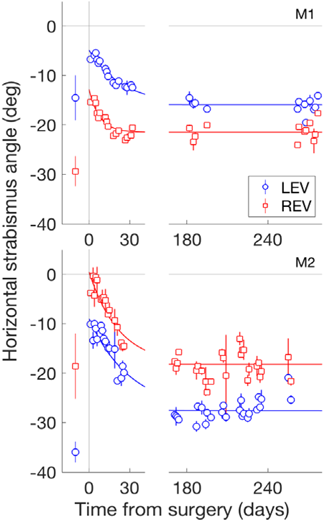
Longitudinal changes in horizontal strabismus angle (top – M1, bottom – M2). The x-axis indicates the time after surgery in days. The data points before the ‘0’ on the x-axis represent the average misalignment before surgery. The negative strabismus angle indicates exotropic misalignment. The blue and red colors correspond to the strabismus angle when viewing with the left eye (LEV) and right eye (REV), respectively. The symbols represent average strabismus angle on a single day, and the lines indicate the best-fitting exponential decay function (decay constants: M1, 24.3days – LEV, 7.7days – REV; M2, 30.8days – LEV, 22.3days – REV). Error bars represent ±SD around the mean.

In total, data from 530 burst-tonic motoneurons were collected in the two awake-behaving animals across different time intervals and in the four motor nuclei. Table 1 shows the distribution of cells collected from each nucleus and at each time interval.

**TABLE 1:**
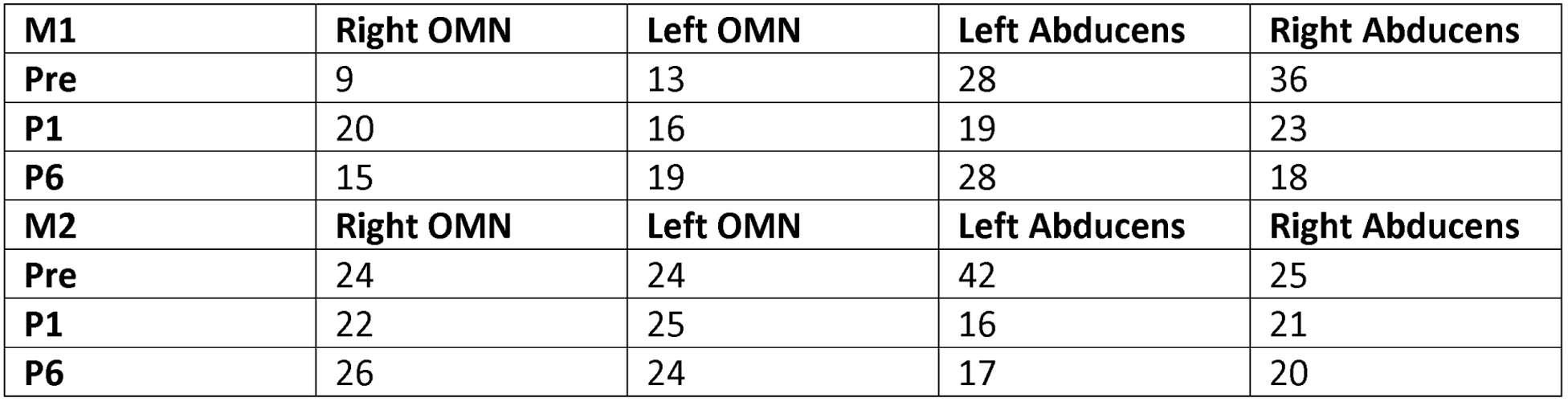
A summary of the number of cells recorded from each oculomotor (OMN) and abducens nucleus in M1 and M2 across each time time-point.

| M1 | Right OMN | Left OMN | Left Abducens | Right Abducens |
| --- | --- | --- | --- | --- |
| Pre | 9 | 13 | 28 | 36 |
| P1 | 20 | 16 | 19 | 23 |
| P6 | 15 | 19 | 28 | 18 |
| M2 | Right OMN | Left OMN | Left Abducens | Right Abducens |
| Pre | 24 | 24 | 42 | 25 |
| P1 | 22 | 25 | 16 | 21 |
| P6 | 26 | 24 | 17 | 20 |

### Changes in K, R and C following surgery

Most studies investigating motoneuron responses have used a first order model (Eq. 1) to characterize response properties. Figure 3 shows two representative cells recorded from the left abducens of monkey M1 (left burst-tonic MNs), one recorded before and the other one-day after surgical treatment. The monkey was able to perform the smooth-pursuit task with normal looking eye movements even the day after surgery and neuronal responses from both these cells are well fit using the first order model with goodness of fit of 0.89 and 0.97 respectively. Average R^2^ value at each time interval were 0.87±0.12; 0.88±0.1 and 0.86±0.14 suggesting that the first-order model was an adequate representation of neuronal responses both before and after surgical treatment of strabismus.

**FIGURE 3:**
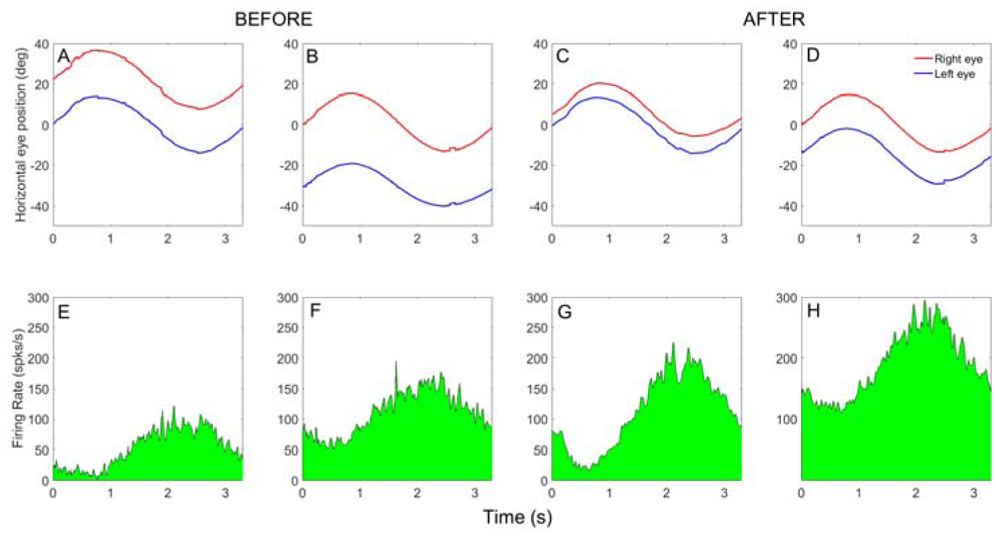
Representative left burst tonic cells recorded from the left abducens nucleus of M1 before and after surgery. The top row shows the average horizontal eye movement traces from multiple cycles of smooth-pursuit. Positive numbers represent rightward eye position and negative numbers represent leftward eye position. Left eye position is shown in blue and right eye position is in red. The bottom row shows the corresponding neural responses (spike density function of firing rates). Panels A, B, E, F are pre-surgery data (K= −2.09 spks/s per deg, R= −1.23 spks/s per deg/s, C=51.4 spks/s, r^2^ = 0.89) and Panels C, D, G, H are from a cell recorded one day after surgical treatment (K= −5.14 spks/s per deg, R= −1.25 spks/s per deg/s, C= 109.1 spks/s, r =0.97).

Figure 4 shows a summary of the average position sensitivity coefficient (K), velocity sensitivity coefficient (R) and baseline firing (C) calculated from equation 1 at each time interval for the four motor nuclei (left and right OMN, left and right Abducens). As described earlier, equation 1 employs the position and velocity of the eye controlled by the muscle to which the neuron projects and smooth-pursuit data from both right eye and left eye viewing conditions were concatenated prior to fitting. Neuronal sensitivity coefficients of neurons projecting to the muscles of the treated eye are shown in panels A, B and C and the coefficients of neurons projecting to the muscles of the untreated eye are shown in panels D, E and F. Fundamentally, the changes in each of the coefficients over the different time intervals were complex and variable across the two monkeys. Overall trends for coefficients, with some exceptions, were to change at P1 but return to close to pre-surgical values by P6.

**FIGURE 4:**
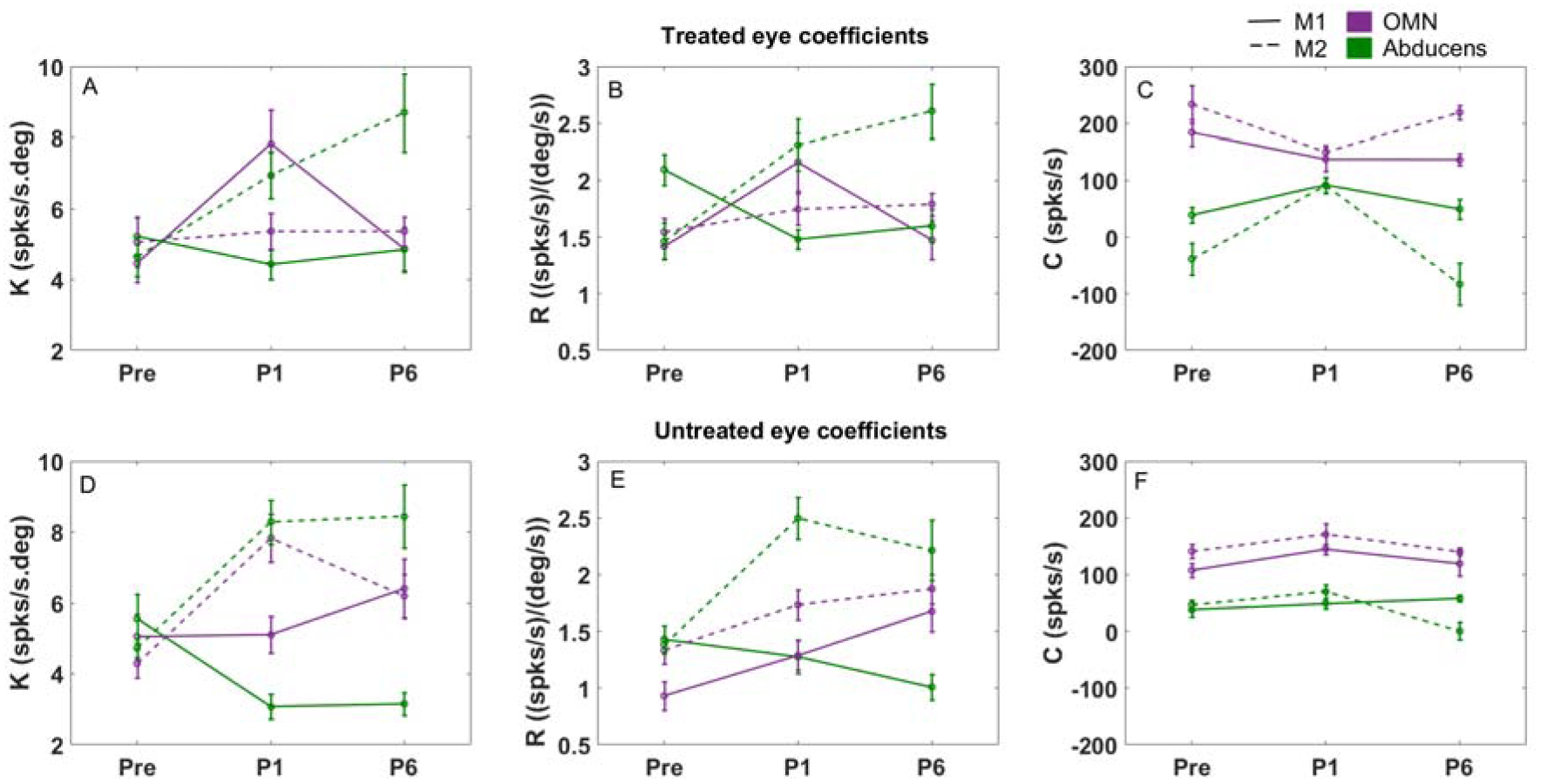
Panels A, B and C show the mean (± standard error) of the position coefficient (K), velocity coefficient (R) and baseline firing (C) at each time interval from motoneurons projecting to muscles in the treated eye in M1 and M2; parameters obtained from cells projecting to muscles in the untreated eye are summarized in panels D, E, and F. The parameters associated with oculomotor neurons are shown in purple and parameters associated with abducens neurons are in green. Solid lines are data associated with monkey M1, and dotted lines are data associated with monkey M2.

The position coefficient ‘K’ obtained from model fitting of the firing rates of Medial Rectus Motoneurons projecting to the treated (left) eye in M1, i.e., RTBT MRMN within left OMN, increased significantly (p=0.01) after surgery (at P1) while the position coefficient (K) associated with the Lateral Rectus Motoneurons (LTBT LRMN within left abducens) projecting to the treated eye showed no significant changes (p=0.64). In M2, the ‘K’ values associated with MRMN or LRMN projecting to the treated right eye (LTBT MRMN within right OMN and RTBT LRMN within right abducens) at P1 showed no significant changes compared to pre-surgical values. ‘K’ values at P6 for both animals were similar to pre-surgical values except for the M2 abducens cells.

The ‘R’ coefficient is related to the velocity of eye movements and is likely to reflect disruption in eye movement dynamics rather than static misalignment. In the treated (left) eye of monkey M1, ‘R’ values decreased for LTBT abducens cells and increased for RTBT OMN cells at P1 but reverted to pre-surgical values at P6 for the OMN cells but not the abducens cells. In the treated (right) eye of M2, ‘R’ values of LTBT OMN cells showed no significant changes at P1 or P6 while ‘R’ values of abducens cells increased after surgical treatment at P1 and P6. In our previously published study describing the eye movement data from these same animals before and after surgical treatment, eye alignment changes were large and obvious but eye velocity changes (i.e., saccade peak velocity and smooth-pursuit gains) were small, idiosyncratic and inconsistent across the animals and directions of eye movement (Pullela et al. 2016). Correlating the dynamic changes in eye movements with the changes in the ‘R’ coefficient was therefore challenging and not pursued in detail.

The coefficient ‘C’ represents the firing rate of the cell when the animal is fixating straight ahead. In general, prior to surgery, the ‘C’ values of MRMN in both the treated and the untreated eyes in our exotropic monkeys were higher than the ‘C’ values obtained from the LRMN and also higher than reported values in normal monkeys of ^~^100 spks/s (Mays and Porter 1984,VanHorn and Cullen 2009). One interpretation of the difference in ‘C’ value between normal and exotropic monkeys is that muscle length adaptation changes the set-point of the eye towards a more abducted location when compared to the normal animal and therefore viewing straight-ahead would involve a relaxation of the LR and contraction of the MR from this new set-point resulting in an increase in ‘C’ values of the MRMN and reduction in ‘C’ values of the LRMN compared to the normal animal.

In the treated eye in M1, changes in C values were not significant at the various time points. In M2, at P1 there was a reduction in ‘C’ values of MRMN (p = 0.02) and increase in ‘C’ values of LRMN (p= 0.002) and these values reverted towards pre-surgical values by P6.

In general, changes in coefficients of cells projecting to muscles of the untreated eye showed similar complex changes. In some cases, the trends were similar to those seen in cells of the treated eye muscles (e.g., lower C values from the left and right abducens MN at P6 compared to P1 in M2) while in other cases, the trends were different (eg, the significant decrease in the C value of the MRMNs within the right OMN compared to no changes in the C value of the MRMN within the left OMN of M2).

### Changes in neuronal drive following surgery

Our goal was to estimate the longitudinal change in neural signals that are driving the covered eye to be deviated at pre- and post-treatment time points. Therefore, to get a handle on the total neural activity that is determining the position of the deviated eye during fixation (state of strabismus), we calculated a Neuronal Drive (ND) to each EOM of the deviated eye (estimated from the population activity within each of the OMN and abducens nuclei - equation 2). If there were no changes in ND across the time intervals, then the longitudinal changes in strabismus angle could be attributed to muscle remodeling factors only. On the other hand, central neural adaptation would manifest as changes in neuronal drive to EOM across the different time points.

Figure 5 shows the estimated average NDs over each time interval to each of the EOMs of the deviated eye while the fellow eye is fixating straight ahead. Panels 5A, C show ND estimates to muscles of the treated eye when the untreated eye is viewing. In general, the NDs from OMN (purple bars) are lower than the abducens NDs (green bars), which could be an indication of the neural basis for exotropia in these monkeys. In monkey M1, in the immediate time period after surgery (at P1 – Panel A), the neuronal drive to the MR is reduced significantly (Holm-Sidak post-hoc comparison, p=0.004) compared to pre-surgical ND while the LR neuronal drive remained the same (p=0.11). Such an overall reduction in ND to only the MR would result in the eye being pushed towards a more exotropic state. The post-surgical eye alignment in M1 at P1 when the untreated eye is viewing is therefore the consequence of beneficial EOM treatment (MR resection and LR recession) plus an adaptive neuronal change to MR drive that is effectively attempting to negate the effect of surgery. By P6, NDs to the treated MR had reverted to pre-surgical values and LR drive remained unchanged. Such a combination should ideally result in an overall reduction of exotropia by P6 but the recurrence of large exotropia suggests a possible role of EOM remodeling over the long term negating the effects of treatment.

**FIGURE 5:**
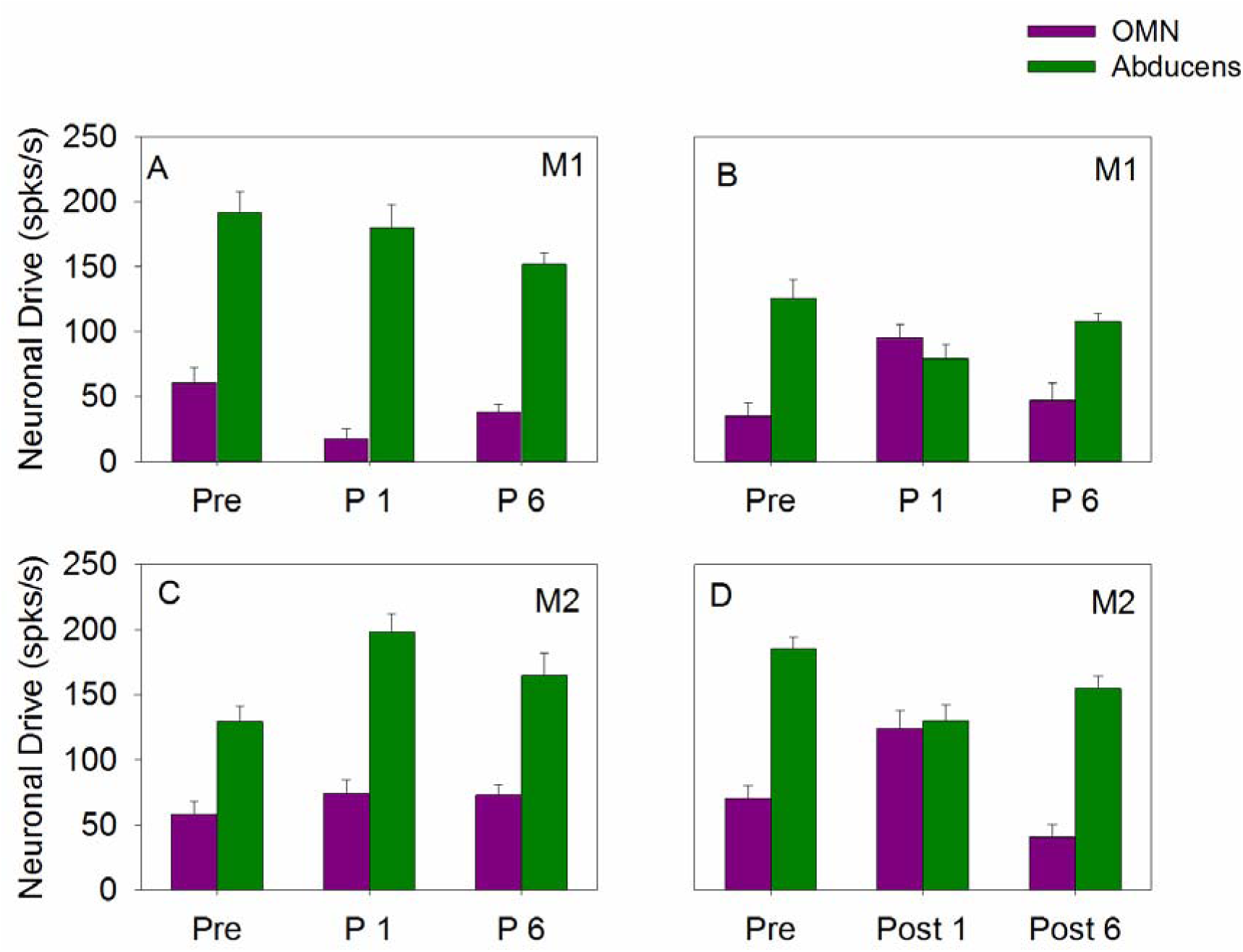
Panels A and B show the mean and standard error of the population neuronal drives (ND) to the EOM of the treated (Panel A) and untreated eye (Panel B) for M1. Panels C and D represent ND to the treated eye and the untreated eye respectively for M2. Drives to the lateral rectus from the abducens nucleus are in green and drives to medial rectus from the oculomotor nucleus are in purple.

The result in M2 at time P1 was different. The treated eye of M2 received an increased ND to the treated LR (p < 0.001) compared to pre-surgery while the ND to the treated MR was unchanged (p= 0.43). An increase to the LR drive is detrimental in that its effect would be to increase exotropia. Therefore, in M2, the post-surgical strabismus angle at P1 observed when viewing with the untreated eye is a result of the benefits of altered muscle contractility due to surgery and a ‘negative’ adaptive neuronal change to the LR. By P6, the NDs to the treated LR reverted to pre-surgical values while the ND to the treated MR remained unchanged. Similar to M1, the recurrence of large angle exotropia suggests a possible role of EOM remodeling.

When the treated eye is forced to view the straight-ahead target (Fig 5B, D), neuronal drive to the muscles of the same *treated* eye must change in comparison to the pre-surgical state to compensate for its altered muscle properties. Therefore neuronal drive to certain muscles of the deviated *untreated* eye will also automatically change due to the anatomical connections between the treated eye abducens nucleus and contralateral (untreated eye) oculomotor nucleus (Buttner-Ennever 2006). This sort of change should not be considered an adaptive response. In M1, the untreated eye received an increase in ND to the MR (p=0.008) and a decrease in ND to the LR (p=0.039) at time P1. While the increase in MR ND may be due to an accompanying change within the abducens of the fellow eye (not an adaptive response), the decrease in ND to the LR is evidence of neuronal adaptation of the untreated eye. The same result was observed in monkey M2 also at time P1. The combined effect results in a smaller strabismus angle after treatment when viewing with the treated eye. Both LR and MR drives reverted towards pre-surgical values by P6.

### Predicted changes in muscle contractility

In addition to the central innervation remodeling, it is likely that some of the final alignment is due to changes in muscle properties. Indirect evidence for muscle remodeling can be gained from the fact that, at P6, strabismus angle and neuronal drives were similar to pre-surgical values although the muscles had been substantially modified by the surgical procedure. Adaptive changes in muscle following EOM surgery or binocular vision disruption have been suggested in other studies also (Hayat et al. 1978, Scott 1994). Although we did not directly measure contractility in this study, we are able to quantitatively estimate the changes in contractility using a simple Hooke’s law based modeling method that uses the eye alignment measures (ϕ) and the recorded neuronal drive (Figure 6).

**FIGURE 6:**
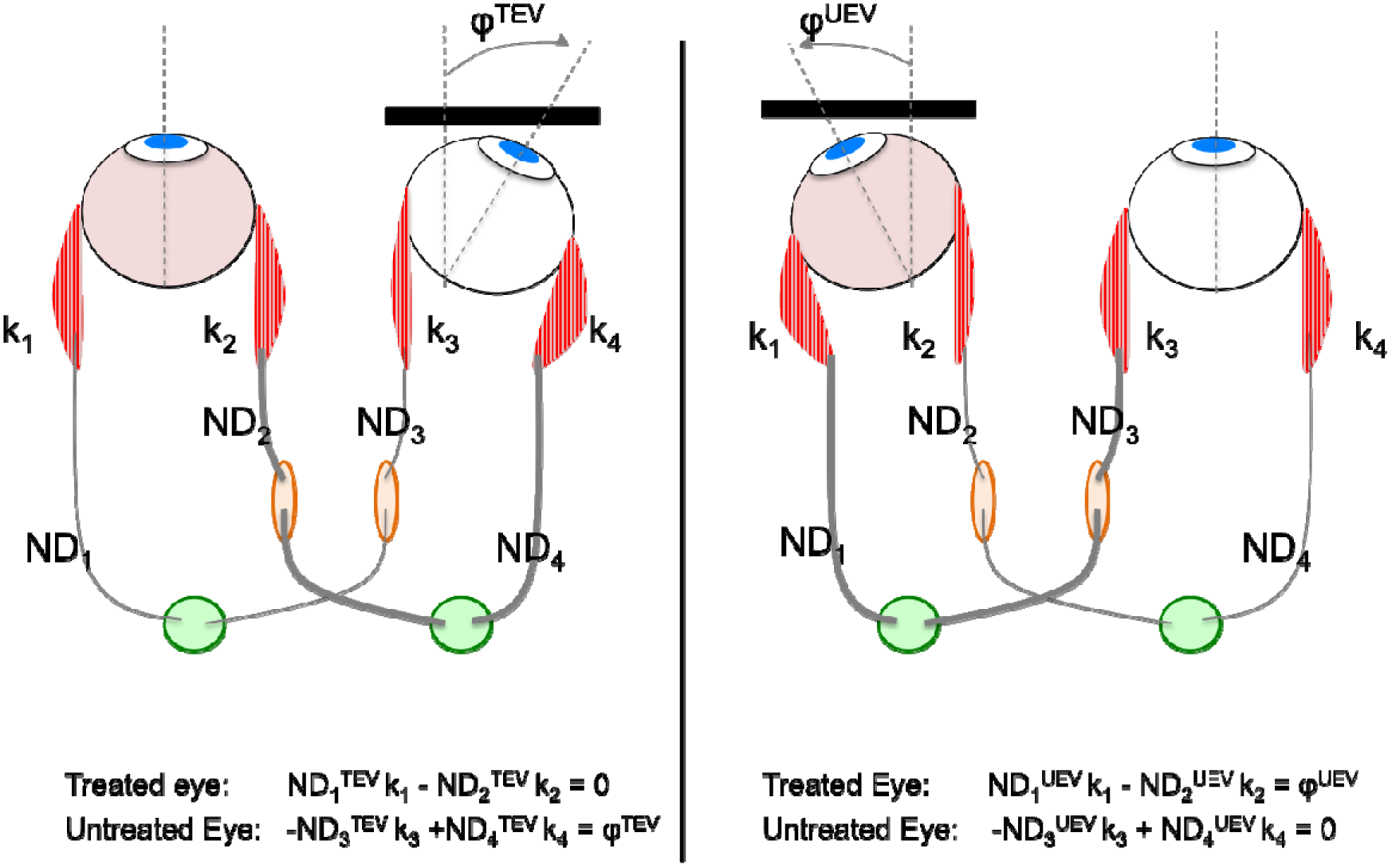
Schematic model for determining muscle contractility. Panel on left develops equations for treated eye viewing conditions (TEV) and panel on right develops equations for untreated eye viewing (UEV) conditions. φ^TEV^ and φ^UEV^ are the positions of the deviated eye during treated eye viewing and untreated eye viewing respectively and is basically the strabismus angle. ND represents the neuronal drive from each nucleus. ‘k1-k4’ represent muscle contractility coefficients and are the parameters to be estimated.

According to Hooke’s law, the force (neuronal drive) needed to extend a spring (EOM) scales linearly with distance (eye position) and is determined by its spring constant (measure of EOM contractility). The equations in Figure 6 describe the Hooke’s law relationship between the ND, EOM spring constants (k1, k2, k3, k4) and the eye positions of the viewing and deviated eyes during either treated eye viewing or untreated eye viewing. The NDs to the muscles of the deviated eye are available from Equation 2 and Figure 4. ND’s to the muscles of the viewing eye (ND1, ND2 during treated eye viewing and ND3, ND4 during untreated eye viewing condition) are simply the ‘C’ term from Equation 1. Since the position of the deviated eye (ϕ^TEV^ and ϕ^UEV^) is basically the strabismus angle and is also a measured quantity, we are able to estimate the contractility coefficients k1 – k4.

The contractility coefficients were obtained using a bootstrapping technique in which 10000 data sets were generated using resampling with replacement. The mean contractility coefficients along with standard errors obtained using this bootstrapping method are shown in figure 7. A one-way ANOVA was used to detect statistical difference among groups and Holm-Sidak post-hoc testing was used for multiple comparisons.

**FIGURE 7:**
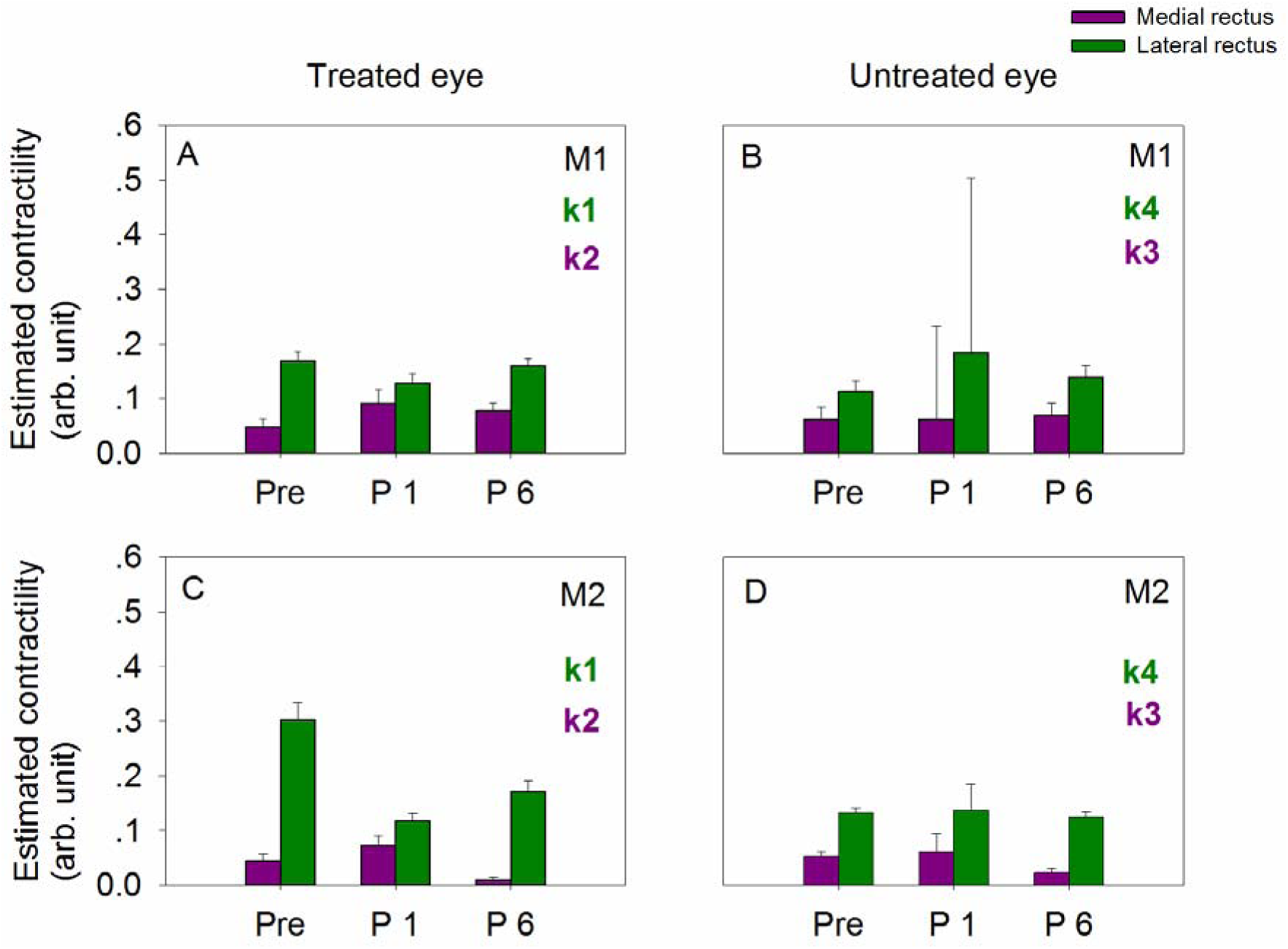
Mean and standard deviation of muscle contractility estimated via the analytical method described in figure 6. Panels A and B represent the MR (k2, k3) and LR (k1, k4) contractility in M1 treated and untreated eye respectively and panels C and D show the MR and LR contractility in M2 treated and untreated eye respectively.

The contractility of the treated LR decreased at P1 with an accompanying increase in the contractility of the treated MR in both M1 and M2. This is to be expected of a LR recess and MR resect surgery. However, an improper adaptive response occurred in treated eye muscles by P6, more pronounced in M2. In both animals, contractility of LR increased and MR decreased compared to P1 indicative of muscle remodeling that contributed to the recurrence of exotropia at P6.

Adaptive changes were also observed in the untreated eye although only in certain muscles. In M1, the contractility of the untreated LR increased significantly at P1 and although decreased by P6 remained higher than the pre-surgical values. In M2 the contractility of the untreated MR did not change much at P1 but decreased to below pre-surgical values by P6. The untreated eye contractility changes in both animals are not beneficial from an alignment perspective as they shift the eye towards exotropia.

It should be noted that our method to assess muscle contractility is indirect at best. Although we expect that this is a reasonable proxy for changes in muscle properties, the most direct method of studying contractility would be to examine specimens of EOM themselves and study muscle contraction properties (Christiansen et al. 2003, Anderson et al. 2006, Anderson et al. 2008).

## Discussion

The main aim of this study was to investigate, longitudinally, the neuronal changes that follow a typical strabismus correction surgery in prism-reared strabismic monkeys. Both monkeys in the study showed large angle exotropia prior to strabismus correction surgery, which decreased by ^~^50-70% immediately after surgery. The strabismus angle underwent rapid change towards increasing exotropia in the first month after correction (0-30 day blocks in Figure 2A, B) and approached pre-surgical levels of exotropia by the end of the P6 period. Recurring exotropia after surgical correction of strabismus is also commonly seen in human subjects and is viewed as a significant clinical problem (Zehavi-Dorin et al. 2016, Lee et al. 2017). Here, we discuss the neuronal and muscle changes that could drive the behavioral changes in alignment following surgery.

### Changes in the neuronal drive from the OMN and Abducens

It is inappropriate to directly correlate changes in sensitivity (coefficients in Eq1) to change in alignment or eye movements because multiple terms in the first-order equation must be considered together to assess neuronal responses during fixation (static – K and C) or eye movements (dynamic – K, R, and C). Further, interpreting the changes in K, R and C across the three time intervals is not a direct reflection of the changes occurring at the neuronal level because the sensitivity coefficients convey the combined effect of the motoneurons and the muscle. For example, when the length of a muscle is changed via resection, the calculated position coefficient K of a cell projecting to a muscle of the deviated eye will change even if the muscle receives the same neuronal command from the motoneurons because the eye is at a different position than before resection. The ND derivation (Eq2) gives a better comparative summary and comprehensive understanding of neural adaptation because it reflects the total output from the brain directed towards a specific extraocular muscle and can be calculated unequivocally for each of the time intervals.

In M1, the ND from the left OMN to the treated MR decreased right after surgery, despite the improvement in the strabismus angle. This works against the aim of the correction surgery as a reduction in ND from the OMN would result in a temporal shift of eye position (towards more exotropia). The ND from the left (treated) abducens did not change at any of the time intervals. This suggests that the immediate improvement in strabismus angle in this monkey is due to the change in effective muscle contractility due to the surgical procedure whose effect was unfortunately partially countered by the reduced neuronal drive to the MR. In monkey M2, the ND from the right OMN to the treated MR remained relatively unchanged following surgery. However, there was unfortunately an accompanying increase in the ND to the treated LR, which will tend to pull the eye temporally (increased exotropia). It was interesting that these changes in neuronal drives were largely reversed by P6.

When the treated eye is fixating, the increase in ND from the untreated OMN to the untreated MR at P1 could, at least partially, be due to an increase in ND within the treated side abducens nucleus because the LR to which the treated abducens projects had been weakened by recession and therefore needs greater innervation (relative to pre-surgery) when fixating a straight-ahead target. Therefore, this increase qualitatively reflects Hering’s law and would not constitute an adaptive response. A post-surgical reduction was also observed in the ND from the abducens to the untreated LR resulting in an improvement in the strabismus while viewing with the treated eye. This is likely to be an adaptive neuronal response whose mechanism is yet unknown. Note however that adaptive muscle changes have been previously reported in the untreated eye of rabbits undergoing EOM surgery (Christiansen et al. 2010) and therefore adaptive neuronal changes to untreated muscle is not entirely unexpected.

In summary, although the misalignment reverted to pre-surgical values in both M1 and M2, it appears that the sequence of neuronal plasticity driving these changes were different in the two animals. Thus, in M1, surgery was rendered ineffective because of reduced ND to MR (‘bad’ plasticity) while in M2, surgery was rendered ineffective because of increased ND to LR (also ‘bad’ plasticity). In both monkeys, the neural plasticity appears to commence immediately after treatment and is largely back to pre-surgical values by 6 months after surgery.

### Combined effects of neuronal plasticity and muscle remodeling

Our analysis of neuronal drives and muscle contractility allows an overall view of what happens as a consequence of strabismus correction surgery. Although both animals showed longitudinal alignment changes that indicate progressive failure of the strabismus correction procedure, the actual neural and muscle changes showed some similarities and some differences between the animals. In the treated eye of both animals, contractility changes following surgery (at P1) were in the appropriate direction (MR contractility increased and LR contractility decreased). However a neural de-adaptive response also commenced (reduction in neural drive to MR in M1 and increase in neural drive to LR in M2) partially offsetting the contractility changes. Although the neural de-adaptive signals were largely gone by P6 (i.e., NDs returned towards pre-surgical values), the contractility changes had also unfortunately reversed. Contractility changes in M2 at P6 appeared to be more severe with a reduction in MR contractility to below pre-surgical values. It was interesting that adaptive neural and muscle changes were observed in the untreated eye of both animals that help to set the strabismus angle when viewing with the treated eye. Specifically changes in ND to LR muscles from the abducens (Fig 5B, D) and the change in MR contractility (Fig 7D) would constitute adaptive responses.

### Other Considerations

Data from this study support a role for unwanted neuronal and muscle plasticity that together results in reversion of strabismus angle. Although the same type of surgery was performed by the same surgeon on the prism-reared experimental monkeys, the plastic changes following surgery were unique to the two monkeys. The variability of plasticity of the neuronal drive and muscle contractility in response to surgery possibly might be the source of the large variability seen in outcomes of strabismus surgery in human patients (Chia et al. 2006). One study reported a success rate of 34% after the initial strabismus correction surgery that rose to 63% after repeated procedures (Habot-Wilner et al. 2012). It is unclear how the brain and the muscles would respond to multiple surgical procedures, and additional primate studies are required to study the effect of multiple procedures on neural and muscle plasticity.

It is also unknown how plasticity might have proceeded if the immediate effect of muscle surgery had been to reduce strabismus angle further than what was currently achieved. It is possible that the nature of plastic changes may have been more helpful if the immediate post-surgical misalignment were small and there was residual binocular vision to ‘lock’ the eyes in an aligned state. Considering the large angle exotropia in our monkeys, operating on EOM of both eyes might have produced better alignment immediately after surgery. However we made the decision to operate on muscles of only one eye to be able to distinguish neural drive changes and muscle changes to a ‘treated’ and ‘untreated’ eye.

In making population estimates of LR neuronal drive, we have used the data from all of the cells when almost certainly part of the recorded population are abducens internuclear neurons (AIN) that project to the contralateral oculomotor nucleus via the medial longitudinal fasciculus (Baker and Highstein 1975, Buttner-Ennever 2006). Other studies that have studied abducens populations have used the same approach (Sylvestre and Cullen 1999,et al. 2014). Fuchs and colleagues (Fuchs et al. 1988)used antidromic activation methods to precisely identify LRMN and AIN and suggested that there was an unique threshold-velocity sensitivity relationship for the LRMN population but not for the AIN population. Cullen et al. used this relationship to construct an upper and lower boundary to classify the population of abducens neurons in their study (Sylvestre and Cullen 2002). Such an approach could not be used in our study due to the horizontal misalignment in our animals. However, it should be noted that Sylvestre and Cullen also showed that the detected AINs do not always encode the monocular position and velocity of the contralateral eye in isolation. Studies have found that groups of abducens neurons identified as AINs and LRMNs behaved similarly during converging eye movements (Mays and Porter 1984) (Gamlin et al. 1989). A more recent study comparing vergence and conjugate sensitivities of normal monkey abducens neurons reported similar findings, suggesting that the response characteristics of AINs and LRMNS are largely similar (Miller et al. 2011). It is unknown whether the same framework is applicable for a strabismic model and further in post-surgical conditions when neuronal drives appears to adapt (Agaoglu et al. 2014). Since we are not unequivocally identifying AIN and LRMN, even if unlikely, the consideration must be left open that post-surgical neuronal plasticity is asymmetric between these neuronal populations.

## Conclusions

In conclusion, we show that both neuronal and muscle plasticity occurs in the aftermath of strabismus correction surgery. Plasticity affects both the treated eye and the untreated eye. Although our sample size is low and therefore inferences must be treated with caution, we suggest that resection is more prone to failure either due to inappropriate neuronal plasticity that drives the eye to exotropia (Monkey M1) or inappropriate muscle plasticity that counters the goal of increased contractility and therefore drives the eye to exotropia (Monkey M2). In any case, most of the neuronal plasticity appears to occur in the immediate aftermath of surgery and so efforts to improve surgical outcomes could be tuned to developing strategies that prevents ‘bad’ plasticity. One such strategy might be to patch the treated eye for a few days after surgery and therefore prevent the brain from initiating an adaptive response driven by vision to the eye whose muscles were manipulated.

